# Compartmental Differences in the Retinal Ganglion Cell Mitochondrial Proteome

**DOI:** 10.1101/2024.05.07.593032

**Authors:** Liam S.C. Lewis, Nikolai P. Skiba, Ying Hao, Howard M. Bomze, Vadim Y. Arshavsky, Romain Cartoni, Sidney M. Gospe

**Author notes:** These authors contributed equally (Romain Cartoni, Sidney M. Gospe).

## Abstract

Among neurons, retinal ganglion cells (RGCs) are uniquely sensitive to mitochondrial dysfunction. The RGC is highly polarized, with a somatodendritic compartment in the inner retina and an axonal compartment projecting to targets in the brain. The drastically dissimilar functions of these compartments implies that mitochondria face different bioenergetic and other physiological demands. We hypothesized that compartmental differences in mitochondrial biology would be reflected by disparities in mitochondrial protein composition. Here, we describe a protocol to isolate intact mitochondria separately from mouse RGC somatodendritic and axonal compartments by immunoprecipitating labeled mitochondria from RGC MitoTag mice. Using mass spectrometry, *471* and *357* proteins were identified in RGC somatodendritic and axonal mitochondrial immunoprecipitates, respectively. We identified 10 mitochondrial proteins exclusively in the somatodendritic compartment and 19 enriched ≥2-fold there, while 3 proteins were exclusively identified and 18 enriched in the axonal compartment. Our observation of compartment-specific enrichment of mitochondrial proteins was validated through immunofluorescence analysis of the localization and relative abundance of superoxide dismutase (*SOD2*), sideroflexin-3 (*SFXN3*) and trifunctional enzyme subunit alpha (*HADHA*) in retina and optic nerve specimens. The identified compartmental differences in RGC mitochondrial composition may provide promising leads for uncovering physiologically relevant pathways amenable to therapeutic intervention for optic neuropathies.

## Introduction

Within eukaryotic cells, mitochondria have diverse physiological roles, including the generation of ATP through oxidative phosphorylation to meet cellular bioenergetic demands, production of reactive oxygen species, buffering of intracellular calcium, and regulation of apoptosis^1–4^. Neurons are particularly reliant on oxidative metabolism, and mitochondria play critical roles in neuronal development, homeostasis, and neurodegeneration^5, 6^. Moreover, the highly compartmentalized nature of neurons is reflected by morphological differences between axonal and dendritic mitochondria, likely a result of divergent mitochondrial dynamics (i.e. fusion and fission) and differential local translation of nuclear-encoded genes functioning in mitochondria^7^. This suggests the existence of mechanisms that regulate mitochondrial function in a manner tailored to the local microenvironments within an individual cell^8–14^. While mitochondrial proteomes have been shown to be distinct between different tissues and cell types^15, 16^, little is known about compartment-specific differences in mitochondrial protein composition in neurons.

Retinal ganglion cells (RGCs), specialized projection neurons of the visual system, are uniquely sensitive to mitochondrial dysfunction. They are prone to degeneration in heritable mitochondrial optic neuropathies that often spare other CNS neurons^17, 18^. Like all projection neurons, RGCs have a highly polarized anatomy^19, 20^. RGC somas localize to the ganglion cell layer (GCL) of the inner retina, while their dendritic arbors extend deeper into the inner plexiform layer (IPL), receiving inputs from presynaptic neurons within their receptive fields. In contrast, the axon of each RGC courses along the innermost retinal nerve fiber layer (RNFL) before converging with other RGC axons and exiting the eye to project via the optic nerve to post-synaptic targets in the brain^21–23^. Due to the extreme morphological polarization between RGC compartments, bioenergetic and other physiological demands differ considerably across the cell^11, 12^. Accordingly, it would be interesting to evaluate whether divergent mitochondrial protein compositions exist between the subcellular compartments in order to meet the specific local needs.

Mitochondrial dysfunction appears to play a key role in the pathogenesis of the most common optic neuropathy, glaucoma—both in primary open-angle glaucoma associated with high intraocular pressure and in the less common variant, normal tension glaucoma^24–27^. In both forms of glaucoma, RGC death and vision loss are the consequence of a degenerative process that begins in a proximal region of the optic nerve called the lamina cribrosa in humans or glial lamina in mice^28–30^. Mitochondrial dysfunction, manifesting as disrupted ATP production and oxidative stress, has been associated with high intraocular pressure in glaucoma patients and in glaucoma mouse models, as well as in the setting of aging, a key risk factor for glaucoma^31–33^. Intriguingly, a recent study in aged glaucomatous mice demonstrated RGC neuroprotection by boosting the level of nicotinamide adenine dinucleotide (NAD^+^), a key cofactor required for mitochondrial ATP production and maintenance of mitochondrial membrane potential^34, 35^. These findings strongly imply a central role for mitochondria in the glaucomatous neurodegeneration of RGCs. However, it is unknown whether axon-specific molecular regulation of mitochondrial function impacts the degeneration of RGCs. The significance of this question is highlighted by a landmark study demonstrating that, whereas deletion of the key pro-apoptosis protein BAX in the DBA/2J glaucoma mouse model completely abrogates RGC soma degeneration, it fails to provide long-term protection to RGC axons in the optic nerve^36, 37^. This result strongly suggests the existence of compartment-specific molecular pathways leading to degeneration in glaucoma^30^.

The goal of this study was to define physiologically relevant, compartment-specific signatures of the RGC mitochondrial proteome. Using an optimized RGC immuno-isolation procedure (‘mitocapture’), we purified GFP-labeled mitochondria from the axonal and somatodendritic compartments of RGC-specific MitoTag mice. Isolated mitochondria were subjected to proteomic analysis aiming to identify proteins differentially expressed in the mitochondria of these compartments, with subsequent validation using immunolocalization techniques.

## Experimental Methods

### Animals

All animal experiments adhered to a protocol approved by the Institutional Animal Care and Use Committee of Duke University. Mice were housed under a 12-hour light-dark cycle with *ad libitum* access to food and water, and all experiments were carried out during the day. *Vglut2-Cre;GFP-OMM^fl/fl^* mice (‘RGC MitoTag mice’) were generated by crossing *Vglut2-Cre* mice^38^ obtained from Jackson Labs (stock no. 028863) with GFP-OMM^fl/fl^ mice^39^ provided by Dr. Thomas Misgeld, Technical University of Munich. C57Bl/6J mice (Jackson Labs stock no. 000664) were used as wild type (WT) controls for all experiments.

### Antibodies

The following antibodies with specified dilutions were used for immunofluorescence (IF) and western blot (WB) analyses: goat polyclonal anti-GFP ([1:1000 WB, 1:750 IF]; abcam, ab5450), rabbit polyclonal anti-HADHA (1:50 IF; abcam, ab54477), rabbit polyclonal anti-SOD2/MnSOD (1:50 IF; abcam, ab13534), mouse monoclonal anti-NDUFS4 (1:1000 WB; Santa Cruz Biotechnology, sc-100567), mouse monoclonal anti-Tuj1 (1:500 IF; Fisher Scientific, MAB11905), mouse polyclonal anti-VDAC (1:1000 WB; EMD Millipore, AB10527), rabbit polyclonal anti-SFXN3 (1:50 IF; Sigma, HPA008028). Secondary antibodies against the appropriate species conjugated to Alexa Fluor 488 (1:1000), Alexa Fluor 568 (1:1000) or Alexa Fluor 680 (1:1000) were purchased from Invitrogen. Cell nuclei were stained using DAPI (Sigma Aldrich). Secondary antibodies against the appropriate species conjugated to IRDye 800CW or 680RD were purchased from Li-Cor for western blot experiments.

### Intravitreal injection of AAV

The plasmid EF1a-mito-dsRED2 was a gift from Thomas Schwarz (Addgene plasmid # 174541; http://n2t.net/addgene:174541; RRID:Addgene_174541). It contains a Flex-Switch-MitoDsRed construct under the transcriptional control of the EF-1α promoter. This construct was packaged into adeno-associated virus serotype 2 (AAV2) by the Duke University School of Medicine Viral Vector Core. 1.25uL of AAV2-Flex-Switch-MitoDsRed viral suspension at a concentration of 3.3×10^12^ vg/mL was injected into the vitreous cavity of anesthetized 3-month-old RGC MitoTag mice, followed by harvesting of optic nerve tissue four weeks later.

### RGC Mitocapture assay

Mitochondria were isolated from freshly collected mouse ocular tissues by adapting a previously described protocol^39, 40^. Adult (P90) RGC MitoTag and WT mice (N=3 each) were euthanized with intraperitoneal administration of a lethal dose of the anesthetic tribromoethanol (Avertin) and transcardially perfused with 1x PBS. Each eye was enucleated and severed at its base from the proximal optic nerve and then placed in 600 µL Isolation Buffer (IB) containing 220 mM Mannitol, 80 mM Sucrose, 10 mM HEPES, 1 mM EDTA, at pH 7.4, and supplemented with 1% essentially fatty acid-free BSA (A7030; Sigma) and 1x protease inhibitors (cOmplete EDTA-free Protease Inhibitor Cocktail; Roche). The calvaria and cortex were removed to expose the optic nerves, which were transected at the optic chiasm and placed into a separate tube containing 600 µL IB. At all stages of the procedure, the samples were kept on ice or at +4°C. Optic nerve tissue was initially minced with Vannas scissors, and retina tissue was dissected and removed from the remaining eye cup. The retina and optic nerve tissues were then separately disrupted in IB with a Dounce glass homogenizer using six up-and-down turns of a Type A pestle. The tissue was then further homogenized with six up- and-down turns of a Type B pestle. The cells were then lysed by nitrogen cavitation using a cell disruption vessel (model 4635; Parr Instrument Company) at 800 PSI, spinning at 80 rpm for 10 minutes. Following a slow pressure release from the nitrogen cavitation vessel, 1x protease inhibitor was added to the resulting tissue homogenate. Nuclei and other cellular debris were removed from the solution by centrifugation at 600 x g for 10 minutes. The supernatant was then transferred to a new tube with IB on ice. Pre-separation filters (#130-041-407; Miltenyi Biotec) were then hydrated with 1 mL of Immunocapture Buffer (ICB) containing 137 mM KCl, 2.5 mM MgCl_2_, 3 mM KH_2_PO_4_, 10 mM HEPES, 1 mM EDTA, at pH 7.4, and supplemented with 1% essentially fatty acid-free BSA and 1x Protease Inhibitor cocktail. The transferred supernatant was then passed through the pre-separation filter and washed with 2 mL ICB. The filtered solution was then centrifuged at 13,000 x g for 3 minutes. The supernatant was removed, and the pellet resuspended in 400 μL ICB.

Subsequently, 100 μL of each sample was removed and set aside as representative “crude mitochondrial fraction.” The remaining 300 μL was diluted to 2 mL total with ICB and combined with 60 μL of superparamagnetic microbeads coated with mouse IgG_1_ antibodies against GFP (#130-091-125; Miltenyi Biotec). These samples were then placed on a rotating wheel at +4°C for 1 hr. LS Columns (#130-042-401; Miltenyi Biotec) were then placed on a magnetic QuadroMACS separator (#130-090-976; Miltenyi Biotec) and equilibrated with 1 mL ICB. To separate the immunocaptured, GFP microbead-coated mitochondria from the remaining cellular material, the solution was then applied to the columns in 1 mL increments, allowing the solution to fully pass through the column until the remaining portion was applied. Importantly, while LS columns are attached to the QuadroMACS separator, bead-antibody complex, and immunoprecipitated material are trapped by the magnetic field. Once the entire sample had passed through the column, each column was washed with 4 x 4 mL ICB. The solution that passed through the column was designated as the flow-through. LS columns were then removed from the QuadroMACS magnetic separator, and GFP-labeled mitochondria were removed from the LS column with 2 x 3 mL washes with ICB, using a plunger to push the remaining immunoprecipitated sample through the column on the last wash. Mitochondria adherent to the microbeads were pelleted by centrifugation at 13,000 x g for 3 min and washed twice with IB without BSA with subsequent centrifugation at 13,000 x g for 3 min between washes. Pellets were stored at −80°C prior to western blot analysis or sample preparation for proteomic analysis.

### Western blotting

For western blot analysis, mitochondrial pellets were resuspended in RIPA buffer containing 50 mM Tris–HCl, pH 8.0, 150 mM NaCl, 0.1 mM EDTA and 1% Triton X-100, 0.25% Nonidet P-40, 0.1% SDS, as well as 1x Protease Inhibitor cocktail on ice for 30 min with frequent vortexing. Samples were mixed with 1x LDS sample buffer (NP0007; Thermo Fisher Scientific) and heated at 70°C for 10 min. SDS-PAGE was performed on the entire sample, followed by transfer to PVDF membrane pre-activated with methanol. Following transfer, the membrane was blocked with a 1:1 mixture of PBS-Tween (0.0125%) and Li-Cor Intercept Blocking Buffer (Li-Cor, catalogue #927-700001), then incubated overnight at +4°C with primary antibody. After washing, the blot was incubated with secondary antibody for 1 hr at room temperature, washed, and imaged using the Li-Cor Odyssey CLx.

### Sample preparation and LC-MS/MS analysis

Protein concentration in the samples was determined with a BCA assay (Pierce BCA Protein Assay Kit; Thermo Fisher Scientific) according to the manufacturer’s instructions, using BSA as the standard. For each sample, 10 μg of total protein was used to prepare peptide mixtures for proteomic profiling. Proteins were cleaved with the trypsin/endoproteinase LysC mixture (Promega, V5072) using the paramagnetic beads-based method^41^. Each digest was dissolved in 12 μl of 1/2/97% (by volume) trifluoroacetic acid/acetonitrile/water solution, and 4 μl were injected into a 5 μm, 300 μm × 5 mm PepMap Neo C18 trap column (Thermo Fisher Scientific) in 1% acetonitrile in water for 3 min at 5 μl/min. Analytical separation of peptides was performed on an EasySpray PepMap 2 μm, 75 μm × 250 mm, C18 column (Thermo Fisher Scientific) over 90 min at a flow rate of 0.3 μl/min at 35°C, using Vanquish Neo UPLC (Thermo Fisher Scientific). The 2-30% mobile phase B gradient was used, where phase A was 0.1% formic acid in water and phase B 0.1% formic acid in acetonitrile. Peptides separated by LC were introduced into the Q Exactive HF Orbitrap mass spectrometer (Thermo Fisher Scientific) using positive electrospray ionization at 2000 V and capillary temperature of 275°C. Data collection was performed in the data-dependent acquisition (DDA) mode with 120,000 resolution (at m/z 200) for MS1 precursor measurements. The MS1 analysis utilized a scan from 375-1450 m/z with a target AGC value of 3.0e6 ions, the RF lens set at 30%, and a maximum injection time of 50 ms. Peptides were selected for MS/MS using charge state filtering (2-5), monoisotopic peak detection, and a dynamic exclusion time of 20 sec with a mass tolerance of 10 ppm. MS/MS analysis was performed using Higher-Energy Collisional Dissociation with a collision energy of 30±5% with detection in the ion trap using a rapid scanning rate, AGC target value of 1.0e5 ions, and maximum injection time of 100 ms.

### Protein identification and quantification

For label-free relative protein quantification, raw mass spectral data files (.raw) were imported into Progenesis QI for Proteomics 4.2 software (Waters Inc.) for duplicate runs alignment of each preparation and peak area calculations. Peptides were identified using Mascot version 2.5.1 (Matrix Science) for searching the UniProt 2019 reviewed mouse database containing 17,008 entrees and supplemented with eGFP amino acid sequence from *Aequorea victoria*. Mascot search parameters were: 10 ppm mass tolerance for precursor ions; 0.025 Da for fragment-ion mass tolerance; one missed cleavage by trypsin; fixed modification was carbamidomethylation of cysteine; variable modification was oxidized methionine and deamidation of asparagine and glutamine. Only proteins identified with 2 or more peptides (peptide false discovery rate (FDR) < 0.5% and protein FDR < 1% calculated using reversed decoy database), were included in the protein quantification analysis. To account for variations in experimental conditions and amounts of protein material in individual LC-MS/MS runs, the integrated peak area for each identified peptide was corrected using the factors calculated by the automatic Progenesis algorithm utilizing the total intensities for all peaks in each run. Values representing protein amounts were calculated as a sum of ion intensities for all identified constituent non-conflicting peptides. Protein abundances were averaged for two technical replicates for each sample, and p-values were calculated for each protein using ANOVA analysis. The abundance ratio between RGC MitoTag and WT control sample was calculated for each protein.

In order to compare somatodendritic and axonal protein abundances, each proteomics dataset was first filtered by removing identified proteins with a Unique Peptide Count < 2, MitoTag/WT ≤ 1.5 and a p-value ≥ 0.05. We then collated the datasets to identify proteins that appeared in at least 3 of 4 biological replicates. After initial filtering of the datasets and protein identification across replicates, MitoTag protein abundance values were normalized to MitoTag GFP values for each dataset and subsequently averaged among biological replicates to create an ‘average GFP-normalized value’ for each protein identified in at least 3 replicates. The ratio between normalized abundances of axonal and somatodendritic proteins was used to identify the relative enrichment of each protein in either compartment. Volcano plots were constructed to highlight the differential abundance of proteins identified in both somatodendritic and axonal compartments. For each identified protein, a t-test was performed to identify significantly enriched proteins in each compartment. Optic nerve to retina (ON/Ret) compartment enrichment ratios and p-values were calculated and plotted on the x and y axes using *log*_2_and −*log*_10_formulas, respectively. An ON/Ret or Ret/ON threshold ≥2 was applied to identify compartmentally-enriched proteins.

### Histological preparation

For immunofluorescence experiments, RGC MitoTag mouse retina and optic nerve tissues were harvested following euthanasia. Enucleated eyes with optic nerves attached were fixed in 4% paraformaldehyde in PBS for 90 min on ice, followed by removal of the anterior segments to produce a posterior eyecup. After initial fixation, optic nerves were excised from the back of each eye. For retinal and optic nerve cross-sections, tissue samples were cryoprotected in sequential 15% and 30% sucrose solutions for 24 hours each. Tissue samples were frozen in Tissue Freezing Medium (Triangle Biomedical), sectioned at 16 μm on a Leica CM1950 cryostat, and placed on glass slides. Sections were hydrated in 1x PBS and blocked in 3% normal donkey serum in PBS with 0.3% Triton X-100. Sections were stained with primary antibodies diluted in blocking buffer overnight at 4°C in a humidity chamber. Sections were subsequently washed and incubated with secondary antibodies for 2 h at room temperature, followed by washing and mounting in Fluoromount G (Invitrogen).

### Quantification of protein abundance by immunofluorescence

To quantify the intensity of colocalized pixels between each protein of interest (SOD2, SFXN3, or HADHA) and GFP, stained retina and optic nerve sections were imaged on a Nikon Eclipse Ti2 inverted confocal microscope using a Nikon A1 confocal scanner system controlled by Nikon NIS-Elements software (Nikon, Tokyo, Japan). Images were acquired using a 40x objective with oil immersion. An 1800 m^2^ region was used to acquire four images across independent retina and optic nerve tissue samples (N=4 of each). The ImageJ “AND” function was used to perform a bitwise comparison between the protein of interest and GFP immunostaining to create a “filtered” image of GFP-colocalized protein, from which the fluorescence intensity was then measured. Filtered fluorescence intensity was averaged across four images for each protein and divided by the average fluorescence intensity of GFP in each region to derive a representation of the abundance of each protein normalized to amount of mitochondrial material.

### Electron microscopy

For ultrastructural analysis, purified mitochondrial pellets from RGC MitoTag retinas and optic nerves obtained via the MitoCapture immunoprecipitation procedure were fixed in a solution of 2% glutaraldehyde and 2% paraformaldehyde in 1x PBS. The specimens were then post-fixed in a solution of 1% osmium tetroxide in 0.1% cacodylate buffer, followed by dehydration with graded ethanol from 30%-100% and propylene oxide. A 1:1 propylene oxide:Embed 812 Resin mixture was infiltrated overnight under a vacuum. Pure Embed 812 Resin was then exchanged on the second day and samples were incubated at 65°C overnight. Ultra-thin sections were then cut at 54-75 nm thickness (Leica EM CU7) and contrast stained with 1% uranyl acetate, 3.5% lead citrate solution. Image acquisition was then performed on a JEM-1400 transmission electron microscope (JEOL) using a Gatan ORIUS (1000) camera.

### Statistics

Statistical analysis was performed using GraphPad Prism software. Specific sample size, statistical test and p-values for each experiment are given in the appropriate figure legends. p-values ≤ 0.05 were considered significant. Standard error bars are shown on graphs.

## Results

### RGC MitoTag mice exhibit expression of GFP-labeled mitochondria throughout RGC somatodendritic and axonal compartments

Previous work has established a protocol for immuno-isolating labeled mitochondria from CNS tissue *in vivo* using a genetically-modified mouse line^39^. The ‘MitoTag’ mouse can be crossed with transgenic mouse lines carrying tissue-specific Cre recombinase drivers to achieve labeling of mitochondria with a cytosol-facing GFP targeted to the outer mitochondrial membrane (‘GFP-OMM’; **Figure 1A**). Mitochondria from the cell type of interest can then be isolated from crude mitochondrial preparations obtained from complex tissues^39, 42^.

**Figure 1.**
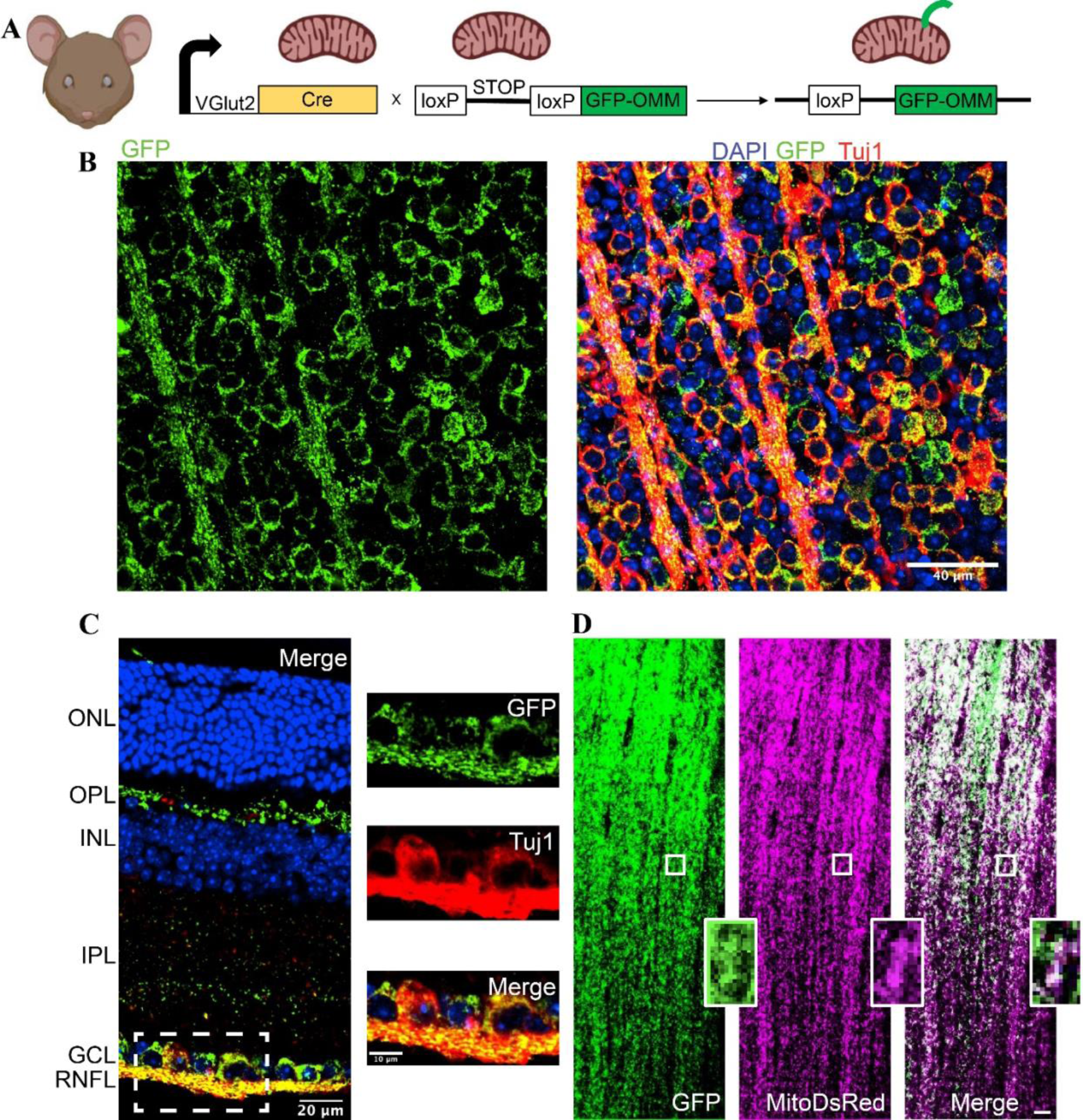
Characterization of the *Vglut2-Cre;GFP-OMM* (RGC MitoTag) mouse. **A.** Schematic of the genetic cross between *Vglut2-Cre* and *GFP-OMM^fl/fl^*mice to generate RGC MitoTag mice in which the STOP codon is excised to allow expression of the GFP-OMM construct in RGCs. **B.** Retinal flat mount from an adult RGC MitoTag mouse showing expression of the GFP-OMM construct (green) in nearly all RGCs (labeled with Tuj1, red). **C.** RGC MitoTag retinal cross-section showing *Vglut2*-Cre-driven expression of GFP-OMM (green) in retinal ganglion cell somas, dendrites, and axons in the inner retina (bottom of image), with expression of the construct also noted in some horizontal cells in the outer plexiform layer. Inset panels show co-localization of GFP-OMM with Tuj1 in RGC somas and axons. Abbreviations: ONL, outer nuclear layer; OPL, outer plexiform layer; INL, inner nuclear layer; IPL, inner plexiform layer; GCL, ganglion cell layer; RNFL, retinal nerve fiber layer. **D.** RGC MitoTag optic nerve longitudinal section showing *Vglut2-Cre*-driven GFP-OMM expression (green) co-localizing with MitoDsRed (magenta) in RGCs transduced by AAV2-MitoDsRed-Flex-Switch. Inset images show co-localization of GFP-OMM and MitoDsRed within a bundle of RGC axons. Scale bar = 20 µm.

To explore the heterogeneity and complexity of compartment-specific RGC mitochondrial populations, we purified RGC mitochondria from the optic nerve containing axonal mitochondria and from retinal tissue enriched in mitochondria localized to the somatodendritic compartments. This was achieved by specifically tagging RGC mitochondria with the GFP-OMM MitoTag. The MitoTag mouse line was bred with the *Vglut2-Cre* line, in which expression of Cre recombinase is driven by the promoter for the vesicular glutamate transporter *Vglut2*^38^. This promoter begins to drive Cre expression in the retina by embryonic day 13 (our unpublished data), and we have previously been shown that it achieves Cre-mediated DNA recombination in >96% of RGCs and in only in a small subset of other retinal neurons but not in glia^43^. Histological characterization of retinal flat mounts from the *Vglut2-Cre;GFP-OMM^fl/fl^*mouse (henceforth denoted ‘RGC MitoTag’ mouse) demonstrated the expected expression of the GFP-OMM construct (green) in both RGC somas and axons co-labeled with Class III β-tubulin, Tuj1 (red, **Figure 1B**). In retinal cross-sections, GFP-OMM expression was observed in the RGC axons located in the RNFL, RGC somas in the GCL and dendrites in the IPL (**Figure 1C**, *left panel*). Higher magnification images highlight GFP labeling of mitochondria in RGC somas and proximal dendrites, as well as in axonal projections co-labeled for Tuj1 (**Figure 1C**, *right panels*). As expected, some GFP-OMM signal was also observed in the outer plexiform layer, corresponding to a subpopulation of horizontal cells expressing *Vglut2-Cre* (**Figure 1C**).

GFP-OMM expression within the optic nerve was very robust (**Figure 1D**). To confirm that this signal arose from RGC axonal mitochondria, we performed intravitreal injections on adult RGC MitoTag mice with adeno-associated virus (AAV2) carrying a MitoDsRed-Flex-Switch. By transducing the somas of RGCs in this way, we could achieve expression of this fluorescent Cre reporter and ultimately label mitochondria throughout all RGC cellular compartments. As expected, along the length of the optic nerve, MitoDsRed co-localized extensively with the *Vglut2-Cre*-dependent GFP-OMM construct expressed endogenously by RGCs in this mouse line (**Figure 1D**, *insets*).

### Compartment-specific isolation of RGC mitochondria

Having confirmed extensive RGC-specific expression of the GFP-OMM construct in RGC MitoTag mice, we next explored whether we could use this mouse line to characterize compartment-specific RGC mitochondrial populations. (**Figure 2A**). Because the exit of the optic nerve from the eye represents a clear anatomic landmark, distal to which only the RGC axonal compartment is present, purifying RGC mitochondria from harvested optic nerve tissue would allow assessment of mitochondria derived exclusively from axons. Conversely, RGC mitochondria purified from retinal tissue would be greatly enriched for those localizing to the somatodendritic compartments. We therefore tested whether tagged RGC mitochondria could be specifically isolated from retinal and optic nerve specimens. Pooled retinas or optic nerves from three mice were homogenized, and differential centrifugation was performed to obtain a crude mitochondrial fraction containing mitochondria and other subcellular organelles from all cells present in the harvested tissue. Incubation of this preparation with paramagnetic nanobeads conjugated to an anti-GFP antibody allowed immunoprecipitation of RGC-derived mitochondria from the crude mixture.

**Figure 2.**
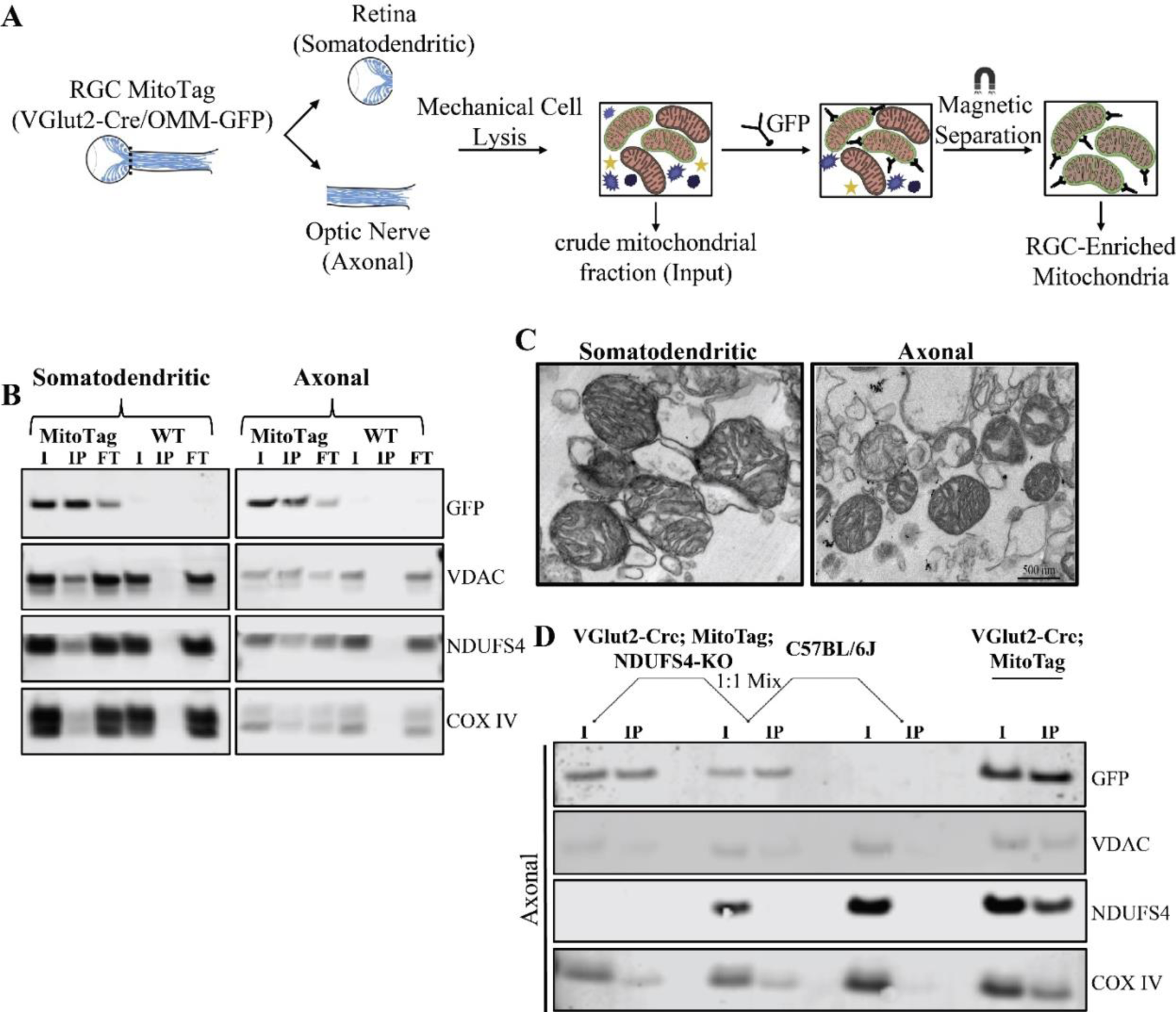
Characterization of the efficiency and specificity of the RGC Mitocapture assay. **A.** Schematic highlighting the workflow for the RGC Mitocapture procedure of obtaining RGC-enriched mitochondrial populations from separate RGC compartments through GFP-based immunopurification. **B.** Representative western blot showing immunoprecipitation of GFP-OMM-tagged mitochondria from RGC somatodendritic and axonal compartments of RGC MitoTAG mice compared to control WT mice lacking the tag. Native mitochondrial proteins VDAC, NDUFS4, and COX4 co-precipitated in RGC MitoTAG but not WT samples. I, input; IP, immunoprecipitate; FT, flow through. **C.** Representative transmission electron micrographs of isolated mitochondria from RGC somatodendritic and axonal compartments. **D.** Western blot from a mixing experiment assessing the specificity of the RGC mitocapture assay. Crude mitochondrial fractions from the optic nerves of *Vglut2-Cre;GFP-OMM^fl/fl^*;*NDUFS4^fl/fl^*and C57BL/6J (WT) mice were mixed at a 1:1 ratio and immunoprecipitated with an anti-GFP antibody. Input fraction from RGC MitoTag optic nerve was used as a positive control.

We used western blot analysis to assess the efficiency and specificity of our purification procedure (**Figure 2B**). To control for nonspecific binding of cellular material to the beads, the purification was also performed on retinal and optic nerve tissue obtained from WT C57Bl/6J mice. The GFP-OMM construct was efficiently immunoprecipitated from the input of both the somatodendritic and axonal RGC MitoTag preparations, along with three other mitochondrial marker proteins: the voltage-dependent anion channel (VDAC), which localizes to the outer mitochondrial membrane, and subunits of respiratory complex I (NDUFS4) and complex IV (COX4), both localized to the inner mitochondrial membrane (**Figure 2B**). As expected, minor fractions of these three proteins immunoprecipitated, as greater proportions derive from unlabeled, non-RGC mitochondria. Importantly, control samples from WT ocular tissues showed negligible precipitation of mitochondrial proteins, indicating low levels of their non-specific binding to the beads. To determine whether purified mitochondria remained intact, we examined the pellets from the immunoprecipitations using transmission electron microscopy and found that the majority of immuno-isolated membranous structures were indeed mitochondria with well-preserved ultrastructure (**Figure 2C**). Additional membranous structures were also observed, likely representing the endoplasmic reticulum, which is known to establish contact sites with mitochondria^44, 45^.

A concern with whole-organelle isolation is that mitochondria from other cell types within a specimen may be recovered nonspecifically alongside RGC mitochondria via clumping during the organelle isolation procedure^46–48^. To assess the extent to which our mitocapture procedure may be affected by such clumping, we performed a mixing experiment to determine qualitatively the extent of non-specific pulldown of unlabeled mitochondria. We generated *Vglut2-Cre;GFP-OMM^fl/fl^;ndufs4^-/-^*mice, in which the GFP-OMM construct is expressed in RGCs of mice with a global deletion of the complex I accessory subunit NDUFS4. Optic nerves obtained from these mice as well as from WT mice and RGC MitoTag mice with intact NDUFS4 were processed to derive mitochondrial samples. Samples from *Vglut2-Cre;GFP-OMM^fl/fl^;ndufs4^-/-^*mice and WT mice were combined at a 1:1 ratio (‘Mixed’ sample), and anti-GFP immunoprecipitation was performed separately on *Vglut2-Cre;GFP-OMM^fl/fl^;ndufs4^-/-^*, Mixed, WT and regular RGC MitoTag samples. Western blot analysis demonstrated a high level of specificity of the RGC Mitocapture assay (**Figure 2D**). When mitochondria tagged with GFP-OMM but lacking NDUFS4 were mixed with WT (non-tagged) mitochondria, no NDUFS4 protein was co-precipitated with the GFP-OMM-positive, NDUFS4-negative mitochondria. This indicates that an overwhelming majority of mitochondrial proteins recovered in the RGC Mitocapture assay were indeed derived from the cell type of interest.

### Compartment-specific heterogeneity of RGC mitochondrial proteins

Having confirmed the efficiency and specificity of the RGC Mitocapture procedure, we characterized mitochondrial proteomes of RGC somatodendritic and axonal compartments using liquid chromatography-mass spectrometry (LC-MS/MS). The analysis was performed on four biological replicates, each with six retinas or optic nerves pooled from three RGC MitoTag mice or WT controls. The data summarized in **Figure 3A** and **Supplementary Tables S1 and S2** include proteins identified from at least 2 unique peptides with p-value ≤ 0.05 whose abundance in RGC MitoTag preparations exceeded the abundance in WT controls by ≥1.5. Based on these criteria, 471 somatodendritic and 357 axonal proteins were identified in at least 3 of 4 biological replicates. Among these proteins, 282 were identified in both compartments, 189 in the somatodendritic compartment and 75 in the axonal compartment (**Supplementary Tables S3, S4, S5**).

**Figure 3.**
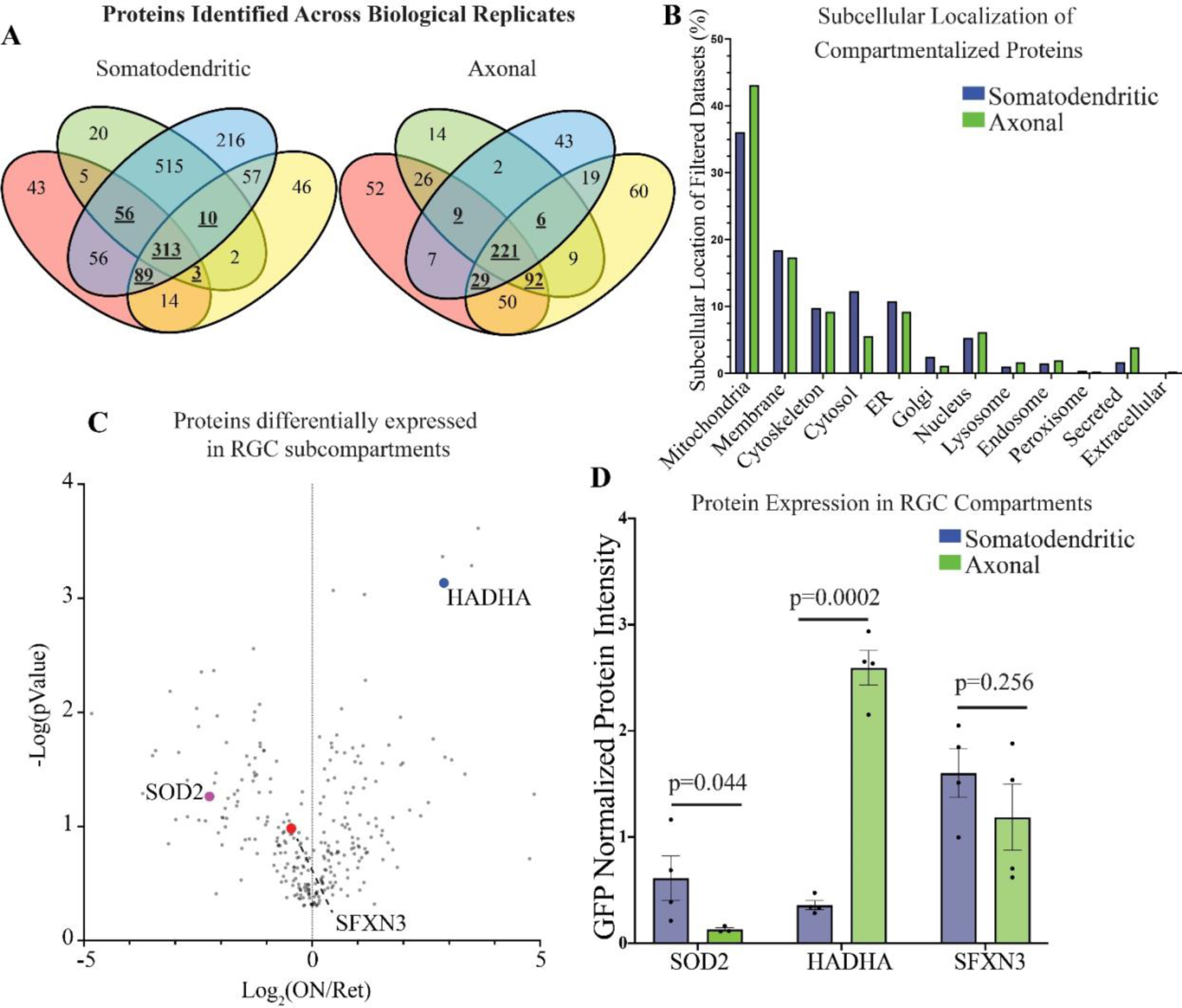
Proteomic profiling of mitochondria isolated from RGC somatodendritic and axonal compartments. **A.** Venn diagrams depicting proteins identified in 4 biological replicates for each compartment. Proteins identified in at least 3 of the 4 replicates are bolded and underlined. **B.** Assigned subcellular localizations of proteins identified in each compartment **C.** Volcano plot showing the relative abundance of proteins identified in both RGC somatodendritic (Ret) and axonal (ON) compartments. Proteins to the left of the vertical line are more abundant in the somatodendritic compartment and those to the right are more abundant in the axonal compartment. Colored dots represent proteins that underwent immunohistochemical validation: SOD2 (purple), SFXN3 (red), and HADHA (blue). **D.** Comparison of GFP-normalized protein abundance for SOD2, SFXN3, and HADHA between the somatodendritic and axonal compartments. A two tailed t-test was performed for each comparison. The GFP-normalized protein abundance for each replicate is depicted as an individual data point.

Using Mitocarta 3.0^49^ and Uniprot ID Mapping^50^, we determined that 36% of all proteins identified in the somatodendritic compartment and 43% of proteins identified in the axonal compartment are known mitochondrial proteins, making them the most highly represented population in these preparations (**Figure 3B**). Additionally, 10% of somatodendritic and 9% of axonal proteins were identified as ER-localized proteins, as expected from the known physical contacts between ER and mitochondria^44^. It is conceivable that at least some of the other identified proteins may have multiple subcellular localizations or were similarly precipitated due to physical contacts between tagged mitochondria and other cellular structures.

For proteins identified in both somatodendritic and axonal compartments, a volcano plot was constructed to compare their relative abundances (**Figure 3C**). To account for any differences in the total amount of mitochondrial material present in each preparation, the total ion intensity of all peptides representing each protein was normalized to that of GFP. The GFP-normalized values were averaged across all biological replicates and the ratios between the two compartments were calculated for each protein. We identified a total of 19 and 18 mitochondrial proteins that were differentially enriched by >2.0-fold in the somatodendritic and axonal compartments, respectively (**Tables I and II**). In addition, 13 mitochondrial proteins were identified in at least 3 of 4 replicates in one compartment but not in a single replicate from the opposing compartment (**Table III**).

**Table I.** Mitochondrial Proteins Enriched in the Somatodendritic Compartment.

| **Protein Name** | **Gene Abbreviation** | **Ret/ON** |
| --- | --- | --- |
| Creatine Kinase b-type | Ckb | 7.60 |
| Superoxide Dismutase [Mn] | Sod2 | 4.69 |
| Cytochrome C Oxidase subunit 4 isoform 1 | Cox4i1 | 4.16 |
| Peroxiredoxin-5 | Prdx5 | 3.93 |
| ATP Synthase Subunit O | Atp5p0 | 3.58 |
| Citrate Synthase | Cs | 3.15 |
| NADH dehydrogenase [ubiquinone] 1 alpha subcomplex subunit 10 | Ndufa10 | 3.10 |
| NADH dehydrogenase [ubiquinone] iron-sulfur protein 7 | Ndufs7 | 2.93 |
| NADH dehydrogenase [ubiquinone] iron-sulfur protein 3 | Ndufs3 | 2.90 |
| Mycophenolic acid acyl-glucuronide esterase | Abhd10 | 2.76 |
| Elongation Factor Tu | Tufm | 2.70 |
| NAD Kinase 2 | Nadk2 | 2.59 |
| Leucine-rich PPR motif-containing protein | Lrpprc | 2.42 |
| 3-hydroxyisobutyrate dehydrogenase | Hibadh | 2.39 |
| Mitochondrial import receptor subunit TOM70 | Tomm70 | 2.317 |
| Mitochondrial-processing peptidase subunit alpha | Pmpca | 2.30 |
| ATP synthase F(0) complex subunit B1 | Atp5pb | 2.20 |
| Prohibitin-2 | Phb2 | 2.08 |
| Glutaminase Kidney Isoform | Gls | 2.03 |

19 mitochondrial proteins enriched by >2.0-fold in RGC somatodendritic mitochondria. Ret/ON values are the ratios of GFP-normalized protein abundances in these compartments averaged across replicates.

**Table II.**
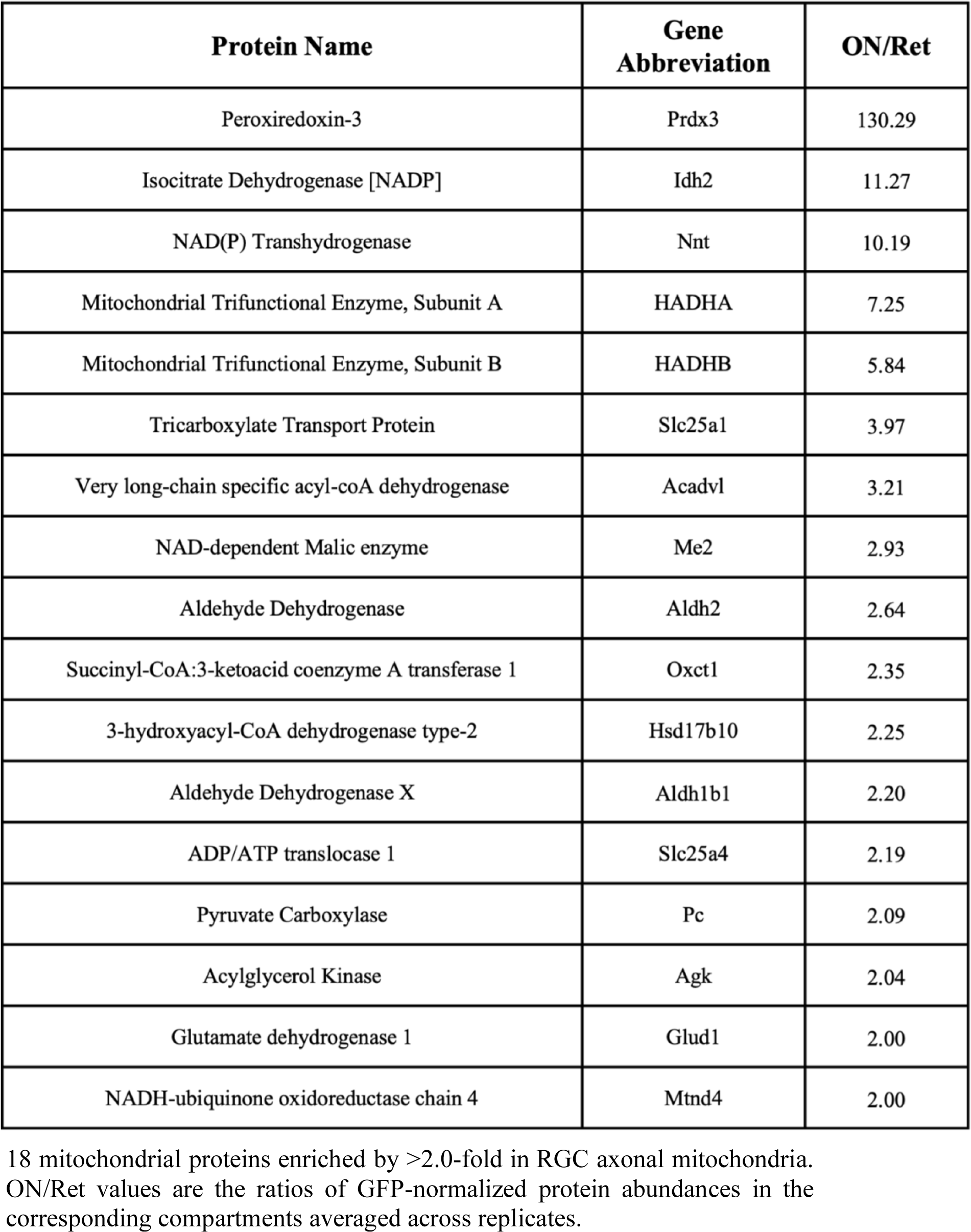
Mitochondrial Proteins Enriched in the Axonal Compartment.

**Table III.** Mitochondrial Proteins Identified in a Single Compartment.

| **Protein Name** | **Gene Abbreviation** | **Compartment Identified In** |
| --- | --- | --- |
| ATP Synthase, subunit D | Atp5d | Somatodendritic |
| Hexokinase-2 | HK-2 | Somatodendritic |
| Tyrosine-tRNA ligase, mitochondrial | Yars2 | Somatodendritic |
| 3-hydroxyisobutyryl-CoA hydrolase, mitochondrial | <u>Hibch</u> | Somatodendritic |
| Mitochondrial import inner membrane translocase subunit TIM50 | Timm50 | Somatodendritic |
| Succinate dehydrogenase [ubiquinone] iron-sulfur subunit, mitochondrial | <u>Sdhb</u> | Somatodendritic |
| Mitochondrial Rho GTPase 2 | Rhot2 | Somatodendritic |
| Elongation factor G, mitochondrial | Gfm1 | Somatodendritic |
| Thioredoxin-related transmembrane protein 1 | Tmx1 | Somatodendritic |
| Caseinolytic peptidase B protein homolog | <u>Clpb</u> | Somatodendritic |
| Alpha-aminoadipic semialdehyde dehydrogenase | Aldh7a1 | Axonal |
| Delta-1-pyrroline-5-carboxylate dehydrogenase | Aldh4a1 | Axonal |
| Choline transporter-like Protein 1 | Slc44a1 | Axonal |

13 mitochondrial proteins identified in either the somatodendritic or axonal compartment in at least 3 biological replicates.

We next sorted proteins confidently identified in either the somatodendritic or axonal compartment according to their reported functions (**Table IV**). Notably, two groups of proteins enriched in the somatodendritic compartment are components of the electron transport chain and those related to antioxidant defenses. Among proteins enriched in the axonal compartment is a group related to fatty acid metabolism.

**Table IV.** Functional Analysis of Compartment-Enriched Mitochondrial Proteins.

|  | Proteins Enriched in Somatodendritic |  | Proteins Enriched in Axonal |  |
| --- | --- | --- | --- | --- |
|  | # Proteins Identified | Proteins Identified (Gene Abbreviation) | # Proteins Identified | Proteins Identified (Gene Abbreviation) |
| Electron Transport Chain | 8 | Ndufa10, Ndufs7, Cox4i1, Ndufs3, Atp5pb, Atp5po, <u>Sdhb</u>, **Atp5d** | 1 | Mtnd4 |
| Lipid Metabolism | 0 | - | 5 | <u>Hadha</u>, <u>Hadhb</u>, <u>Acadvl</u>, <u>Agk</u>, Hsd17b10 |
| ROS Management | 4 | Sod2, Prdx5, Abhd10, **Tmx1** | 1 | Prdx3 |
| Other Metabolic Proteins | 7 | Cs, <u>Ckb</u>, <u>Gls</u>, <u>Hibadh</u>, **Hibch**, NADK2, **HK2** | 13 | Idh2, Pc, Aldh1b1, Oxct1, Aldh2, Slc25a4, Me2, Glud1, <u>Nnt</u>, **Aldh4a1**, **Aldh7a1**, **Slc25a1**, **Slc44a1** |
| Other | 10 | <u>Tufm</u>, Tomm70, Pmpca, Phb2, <u>Lrpprc</u>, <u>Clpb</u>, **Gfm1**, **Rhot2**, **Timm50**, **Yars2** | 0 | - |

Functional classification of 29 and 20 mitochondrial proteins identified in the somatodendritic or axonal compartments, respectively

### Histological validation of compartment-specific expression of RGC mitochondrial proteins

In the next set of experiments, we validated some of our proteomic findings using immunofluorescence analysis. We selected three mitochondrial proteins to confirm their compartment-specific enrichment: mitochondrial superoxide dismutase (SOD2, found to be enriched in the somatodendritic compartment); sideroflexin 3 (SFXN3, equally distributed between compartments); and trifunctional enzyme subunit α (HADHA, enriched in the axonal compartment) (**Figure 3D**). We immunolabeled retinal and optic nerve cross-sections from RGC MitoTag mice with antibodies against each protein. In order to limit our analysis to protein expressed by RGCs but not other cell types, we used *ImageJ* to identify pixels containing signal for both GFP and protein of interest to create a GFP-localized mask for each protein^51, 52^. We then quantified the signal intensity for each protein of interest relative to that of GFP-OMM as a measure of the abundance of each protein within RGC mitochondria (**Figure 4A and 4B**). Consistent with the proteomic data, this immunofluorescence analysis showed a statistically significant enrichment of SOD2 in the somatodendritic compartment and HADHA in the axonal compartment, while no statistically significant difference in SFXN3 abundance was observed between the two RGC compartments (**Figure 4C**).

**Figure 4.**
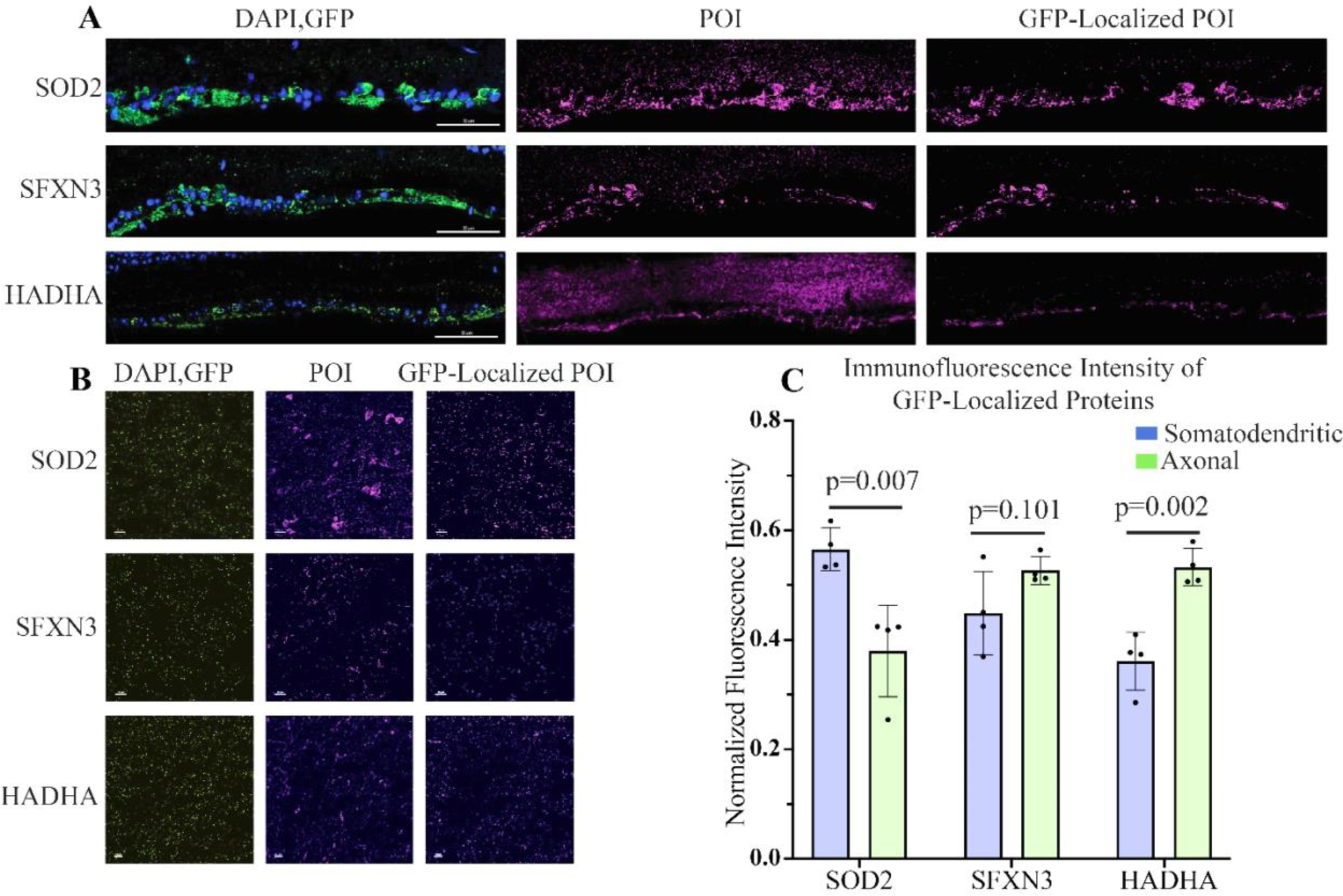
Histological validation of compartment-specific protein abundance in RGC mitochondria. **A,B.** Retinal (A) and optic nerve cross sections (B) from RGC MitoTag mice showing *Vglut2-cre*-dependent expression of GFP-OMM in RGCs (left panels, green), endogenous expression of proteins of interest (POI; middle panels, magenta), and POI expression solely within GFP-positive mitochondria (right panels, magenta). Protein identities are indicated to the left. Scale bars, 50 µm (A) and 20 µm (B). **C.** Comparison of GFP-normalized fluorescence intensities of SOD2, SFXN3 and HADHA in retinal and optic nerve sections from RGC MitoTag mice (N=4 replicates for each tissue). A two tailed t-test was performed for each comparison. The GFP-normalized fluorescence intensities for each replicate are depicted as individual data points.

### Age-related heterogeneity of RGC mitochondrial proteins

Because age is a major risk factor for the development of glaucoma^33^, we proceeded to use the Mitocapture procedure to isolate and characterize mitochondria from young and aged mice. RGC somatodendritic and axonal mitochondria were separately immunopreciptated from 3-month-old and 12-month old RGC MitoTAG mice as well as 3-month-old WT mice as controls (N=3 mice per group for each of two biological replicates). We again used western blot to validate the efficiency and specificity of our isolations. The GFP-OMM construct was immunoprecipitated from the input of both the somatodendritic and axonal RGC MitoTag preparations for both age groups, along with the inner mitochondrial membrane proteins COX4 and NDUFS4 (**Figure 5A**). The failure of COX4 and NDUFS4 to be immunoprecipated from WT control preparations confirmed excellent specificity of the purifications.

**Figure 5.**
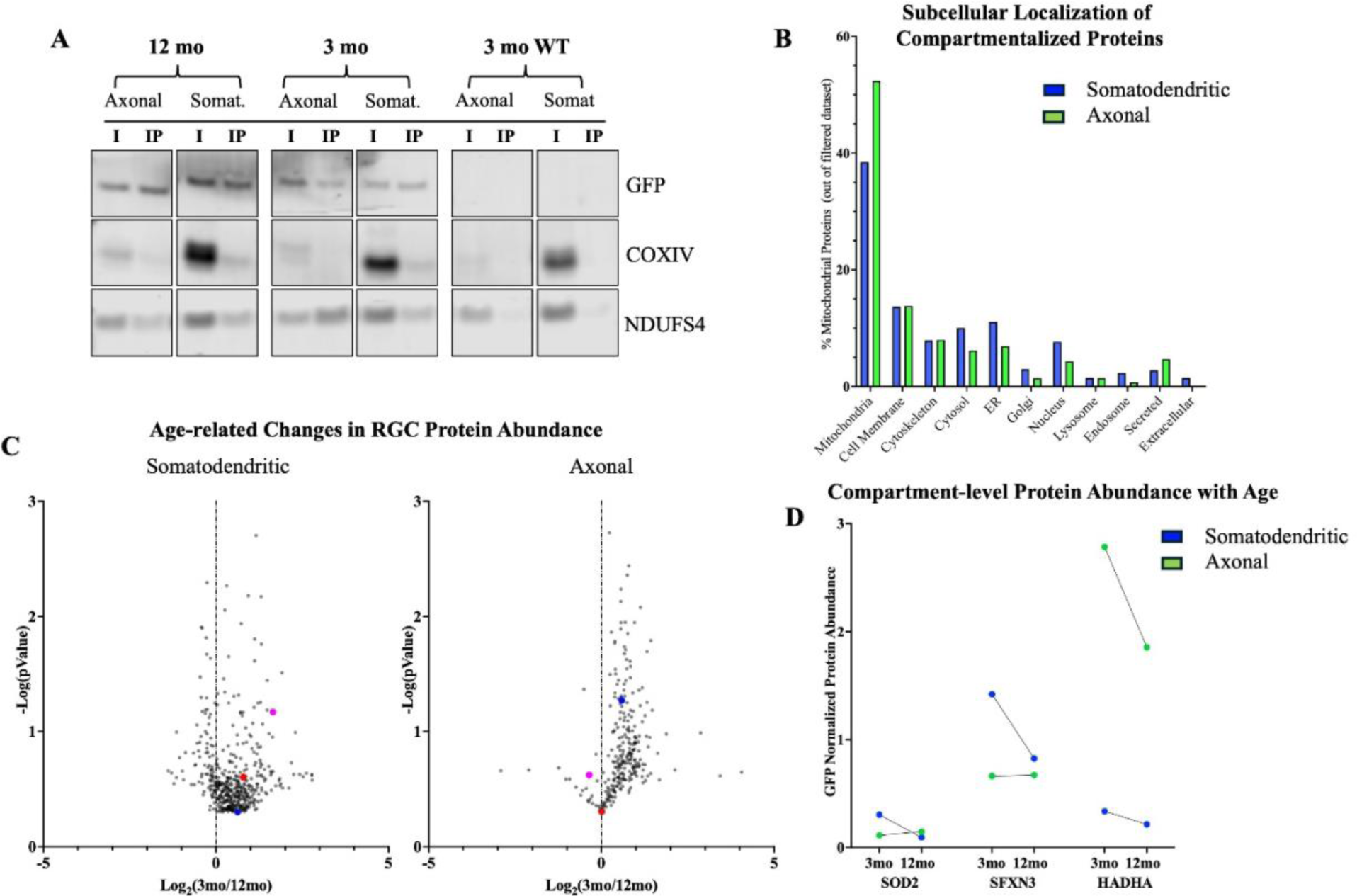
Proteomic profiling of mitochondria isolated from RGC somatodendritic and axonal compartments of young (3 mo) and aged (12 mo) mice. **A.** Representative western blot showing immunoprecipitation of GFP-OMM-tagged mitochondria from RGC somatodendritic and axonal compartments of 3 mo and 12 mo RGC MitoTAG mice compared to control WT mice lacking the tag. Native mitochondrial proteins COX4, and NDUFS4 co-precipitated in RGC MitoTAG but not WT samples. I, input; IP, immunoprecipitate. **B.** Assigned subcellular localization of proteins identified across two biological replicates in each compartment. **C.** Volcano plots showing the relative age-related abundance (3mo/12mo) of proteins identified in both age groups. Proteins to the left of the vertical line are more abundant in 12 mo mice and those to the right are more abundant in 3 mo mice. Colored dots represent proteins of interest: SOD2 (purple), SFXN3 (red), and HADHA (blue). **D.** Comparison of age-related changes in GFP-normalized protein abundance for SOD2, SFXN3, and HADHA for each compartment. Each dot represents an average between two biological replicates.

As before, using LC-MS/MS, we analyzed proteins with at least 2 unique peptides, a p-value ≤ 0.05, and whose abundance in RGC MitoTag preparations exceeded the abundance in WT controls by ≥1.5. Across the two biological replicates and among both age groups, there were 468 and 275 proteins identified in the somatodendritic and axonal compartments, respectively (**Supplementary Tables S6 and S7**). Using the Mitocarta 3.0 and Uniprot ID Mapping protein databases, we determined that 38% of all proteins identified in the somatodendritic compartment and 52% of proteins identified in the axonal compartment are known mitochondrial proteins (**Figure 5B**). For proteins identified in the somatodendritic and/or axonal compartments in both replicates for the 3- and 12-month-old mice, a volcano plot was constructed to compare the relative abundances at the two ages (**Figure 5C**). As before, normalization of the total ion intensity of all peptides for each protein to that of GFP was performed to account for differences in total mitochondrial material present in each preparation. The GFP-normalized values were averaged across two biological replicates and the ratios between the two age groups were calculated for each protein. We determined that the relative abundances of SOD2, SFXN3, and HADHA were all decreased in the mitochondria of the somatodendritic compartment of the 12-month-old mice relative to their counterparts in the 3-month-old mice, while that of HADHA was also decreased in the axonal compartment (**Figure 5D**). Interestingly, this was part of an overall trend of decreased protein abundance in RGC mitochondria immunoprecipated from older mice, with 80% of all proteins less abundant in somatodendritic mitochondria and 91% less abundant in axonal mitochondria at 12 months compared to 3 months. The age-related changes in the abundance and activities of specific proteins are likely to be important in explaining the progression of optic neuropathies like glaucoma and will each require experimental validation. However, our general observation of decreased mitochondrial protein abundance at 12 months may be consistent with the known age-related global decline in mitochondrial function^53–55^.

## Discussion

In this study, we report a novel approach to isolate and characterize intact mitochondria derived from distinct subcellular compartments of a specific cell type. Using LC-MS/MS, we identified mitochondrial proteins enriched in either the somatodendritic or the axonal compartments of RGCs. Remarkably, many of the proteins with compartment-specific enrichment belong to groups with distinct mitochondrial activities, which suggests the existence of local differences in mitochondrial physiology in these neurons. Our approach should be readily adaptable to the compartment-specific analysis of organelles from any projection neurons obtained from living tissues.

### Methodological considerations

Our approach to achieve compartment-specific purification of mitochondria from a cell type of interest was an adaptation of the strategy of Fecher and colleagues to immuno-isolate cell-specifically tagged mitochondria from tissues^39^. *In vivo* tagging of mitochondria in genetically modified mouse strains allows for highly specific isolation of mitochondria deriving from a specific cell type regardless of the complexity of the tissue. However, while the immuno-isolation step is critical, the quality of the mitochondrial purification is still highly dependent on the earlier, more crude steps of the preparation. In accordance with Fecher et al.^39^, we elected to utilize differential centrifugation in obtaining crude mitochondrial fractions, as opposed to gradient-based protocols^56, 57^. While gradient-based isolation methods, such as those using sucrose or Percoll gradients, can result in greater purity of isolated mitochondria, they force mixed populations of mitochondria into closer proximity and increase the likelihood of adherence/fusion amongst mitochondria originating from different cells^42^. We also employed mechanical rather than detergent-based tissue homogenization in order to better preserve mitochondrial integrity to avoid losing mitochondrial proteins from the sample.

Our data indicate that we successfully purified intact mitochondria from RGCs, as normal morphology was observed with electron microscopy and efficient recovery of both inner and outer mitochondrial membrane proteins was demonstrated by western blotting. We adjusted our protocol to optimize the specificity of the immunoprecipitations, although this level of stringency did come at the cost of some efficiency, as the presence of GFP in the flow-through indicates that a fraction of GFP-tagged RGC mitochondria failed to immunoprecipitate from both compartments (**Figure 2B**). Using a mixing experiment, we further showed that the Mitocapture procedure avoids significant nonspecific clumping of mitochondria deriving from other cell types. Thus, we conclude that in our approach the intact mitochondria present in the immunoprecipitation overwhelmingly derive from the cell type of interest.

After completing the LC-MS/MS analysis of immuno-isolated mitochondria, our goal was to compare the abundance of proteins recovered from the somatodendritic and axonal compartments, in order to assess for differential compartment-specific regulation of mitochondrial protein composition. As in other mitochondrial proteomic analyses, we identified a large number of ‘non-mitochondrial’ proteins in immuno-isolated mitochondria from both compartments. While in most cases this can be explained by physical attachment between mitochondria and other cellular membranes and the cytoskeleton, it is also conceivable that some of them reflect previously unknown transient mitochondrial interactions^58^.

Two important caveats of our work must be discussed. First is the fact that the most proximal region of RGC axons exists in the innermost surface of the retina (the RNFL), and therefore some RGC axonal mitochondria will necessarily be isolated alongside somatodendritic mitochondria from retinal specimens. This is illustrated in Figure 4A, in which RGC expression of the axon-enriched protein HADHA appears to be limited almost exclusively to the RNFL rather than strongly labeling the RGC somas and dendrites like SOD2 and SFNX3. It is unavoidable, therefore, that the presence of the RNFL leads to some ‘contamination’ of the somatodendritic mitochondrial preparation with some axonal mitochondria, potentially diluting the degree of compartment-specific differences between the mitochondrial proteomes.

The second caveat is the observed expression of the GFP-OMM tag by horizontal cells in the OPL of the RGC MitoTag mouse retina (**Figure 1**). Because the *Vglut2*-driven Cre is not entirely specific to RGCs, a mitochondrial purification from whole retina will inevitably recover some tagged mitochondria derived from horizontal cells. We previously showed that ∼10% of horizontal cells demonstrate Cre-mediated genetic recombination in *Vglut2-Cre* mice, compared to >96% of RGCs^43^. Based on the literature, there are approximately 45,000 RGCs and 18,000 horizontal cells per mouse retina^59, 60^. If 10% of horizontal cells express Cre recombinase in RGC MitoTag mice, then they should be outnumbered by RGCs by approximately 24:1, which makes their contribution unlikely to significantly obscure compartment-specific mitochondrial proteome differences within RGCs. Nevertheless, due to both of the caveats we raised, it is particularly important that interesting differences identified in our proteomic analyses be validated using immunohistochemical techniques.

### Potential biological significance of compartment-specific differences in the protein composition of RGC mitochondria

Our data suggest the possibility of functional compartment-specific differences in the RGC mitochondrial proteome. HADHA, which we found to be enriched in axonal mitochondria, is the α-subunit of the mitochondrial trifunctional protein (MTP)^61^. It forms a heterotetrameric complex with the β-subunit, HADHB, which is similarly enriched in the same compartment. The MTP complex catalyzes the final three steps in the mitochondrial fatty acid β-oxidation pathway (FAO), ultimately producing acetyl-CoA as a substrate for the TCA cycle^62^. Among other mitochondrial proteins enriched in the axonal mitochondria in our analysis are very-long-chain-specific acyl-CoA dehydrogenase and 3-hydroxyacyl CoA dehydrogenase, which catalyze the first step of β-oxidation of very long chain fatty acids and an intermediate step in FAO, respectively. This constellation of proteins enriched in axonal mitochondria raises the possibility that FAO may be of particular importance to axonal physiology, either as an energy source or for generation of acetyl-CoA as a biosynthetic precursor. Notably, MTP deficiency can lead to a rare, autosomal recessive disorder that causes impaired mitochondrial metabolism of long-chain fatty acids^63^. Among the most frequent disease manifestations of MTP deficiency is peripheral neuropathy, which may indicate a particular sensitivity of projection neurons to aberrant FAO^64^. It would be of particular interest to determine whether RGC-specific impairment of FAO leads to optic atrophy in mice, possibly with axonopathy preceding the death of RGC somas in the retina.

Within the somatodendritic compartment, we identified and immunohistochemically validated an enrichment of SOD2, a critical mitochondrial antioxidant protein that converts superoxide radicals to hydrogen peroxide^65^. Another protein enriched in somatodendritic mitochondria is peroxiredoxin-5 (PRDX5), which can reduce hydrogen peroxide to water, among other functions^66, 67^. The abundance of these antioxidant proteins may suggest that the RGC somatodendritic mitochondria are subjected to especially high levels of oxidative stress and/or require particularly efficient antioxidant pathways to scavenge free radicals and avoid oxidative damage. It is notable that a number of mitochondrial optic neuropathies have been associated with RGC oxidative stress^68–73^.

Consistent with a high degree of sensitivity of the RGC soma to oxidative stress, we have previously shown in a mouse model of RGC-specific NDUFS4 deficiency that the onset of RGC soma degeneration appears to begin prior to loss of axons. *In vivo* work has further shown that adeno-associated virus-mediated overexpression of SOD2 attenuated RGC death in rats with high intraocular pressure^74^. SOD2 overexpression was also observed to inhibit apoptosis, reduce oxidative stress, and increase mitochondrial function in immortalized skin fibroblasts from patients with complex I deficiency and in cultured cells treated with the complex I inhibitor rotenone^70, 75^. Thus, a better understanding of antioxidant pathways in RGC somas may be of particular therapeutic relevance to mitochondrial optic neuropathies.

Another notable finding is that PRDX5 is enriched in RGC somatodendritic mitochondria, whereas the related protein PRDX3 is very highly enriched in RGC axonal mitochondria. Among all members of the peroxiredoxin family, PRDX3 and 5 are the only proteins that localize to mitochondria (PRDX3 is exclusively mitochondrial, while PRDX5 can also exist in peroxisomes and the cytosol)^76^. PRDX3 and 5 are hypothesized to act as ROS sensors and peroxide scavengers in environments of high oxidative stress^77, 78^. It is unclear to what extent PRDX3 and 5 are redundant in their functions, but the striking compartmental discrepancy in their abundance within RGC mitochondria is quite intriguing. Genetically modified mouse lines with loss of PRDX3 and PRDX5 have been generated and reportedly have relatively mild signs of systemic mitochondrial dysfunction^67, 79^. It would be interesting to determine whether loss of either protein is associated with early compartment-specific RGC degenerative phenotypes.

It is also striking that we identified a number of components of the electron transport chain, particularly subunits of complex I and ATP synthase, as being enriched in the RGC somatodendritic compartment relative to the axonal compartment. While the relative abundances of these proteins and the relative activity levels of the electron transport chain in the two compartments remain to be validated, it would seem surprising for axonal mitochondria to have a lower ATP-generating capacity than their counterparts in the RGC soma and dendrites. However, recent work in cortical projection neurons has cast doubt on the role of oxidative phosphorylation in providing for the energy needs of axons, potentially indicating that mitochondrial functions other than ATP generation may be of particular importance in neuronal axons^80^.

## Acknowledgements

This work was supported by the BrightFocus Foundation to R.C. and S.M.G., by a grant from the National Eye Institute (R01EY030969) to R.C. and S.M.G., by National Eye Institute Core Grant EY005722 to Duke University, and by an unrestricted grant from Research to Prevent Blindness to the Duke Eye Center.

## Data availability

The mass spectrometry proteomics data have been deposited to the ProteomeXchange Consortium via the PRIDE partner repository with the dataset identifier PXD051899 and 10.6019/PXD051899.

## Supporting Information

- (Table S1) All Axonal Proteins (XLSX)
- (Table S2) All Somatodendritic Proteins (XLSX)
- (Table S3) Proteins from Both Compartments (XLSX)
- (Table S4) Exclusively Axonal Proteins (XLSX)
- (Table S5) Exclusively Somatodendritic Proteins (XLSX)
- (Table S6) Axonal Aged vs. Young (XLSX)
- (Table S7) Somatodendritic Aged vs. Young (XLSX)

